# Daily Patterns of Foraging and Aggressive Behaviors in Great-tailed Grackle (*Quiscallus mexicanus*) at an Urban Patch with Availability or Absence of Resources

**DOI:** 10.1101/2021.06.14.448443

**Authors:** Alejandro Rodrigo, Laurent Ávila-Chauvet, Jonathan Buriticá

## Abstract

The Great-tailed Grackle *(Quiscalus mexicanus)* seems to take advantage of inhospitable environments such as cities. However, it is not yet fully understood how these birds exploit hostile environments to their advantage. Casual observation seems to suggest that this species can obtain resources of biological importance such as food or nesting material from the trash. As a first approach to the problem, we located a patch outside a residential building, in a high-density urban area, where the residents left their trash for pick-up, and a group of wild Great-tailed grackles was identified as regular visitors. In total, 25 days were recorded at the site (November 2017 - January 2018). Events such as foraging, number of subjects present at the foraging site, aggressive behaviors between members of the group, and their relation with the presence or absence of the garbage collector truck were registered. The results show a higher number of grackles at the observation site and a higher frequency of foraging behaviors in the presence of garbage collection than in its absence. In its presence, the distribution of foraging during the day follows a normal distribution. In the absence, the distribution shows more variability towards the day. The highest frequency of interactions occurred between two grackles, yet there were records of up to eight subjects. The highest number of aggressions registered took place in the absence of garbage collection than in its presence. Moreover, the focal subject exhibits fewer agonistic behaviors compared to other members of the group, a result expected if the producer-scrounger game literature is considered. The outcome is explained in terms of deprivation and availability of resources. Finally, we conclude that grackles can exploit hazardous environments such as cities due to the highly social behaviors exhibited during foraging.

## Introduction

The expansion of human settlements like cities has become one of the most common environments that species can find (Evans et al., 2012; Gallo & Fidino, 2018). Urbanization process is associated with alterations in the environment such as structural and physical changes, replacement of natural vegetation, increased temperatures, air and soil pollution, aggravated levels of annual rainfall, among others. All these factors together tend to change the interactions between and within species populations (McKinney, 2002; Sol, Lapiedra, & González-Lagos, 2013).

Although most animals have the ability to modify their behavior in response to variations in the environment, the rapid and drastic changes in cities have provoked a severe negative impact on a considerable number of species such as extinction and displacement (M. T. J. Johnson & Munshi-South, 2017; McKinney, 2002). As a result, cities are considered one of the most extreme environments for animals. However, some species have been able to adapt to these harsh conditions. Invasive species such as rats, mice, pigeons, cockroaches, mosquitoes, and some others have managed to efficiently exploit these environments, which has benefited their settlement (M. T. J. Johnson & Munshi-South, 2017; Sol et al., 2013).

The Great-tailed Grackle (*Quiscalus mexicanus*, hereafter referred to as grackles) is a remarkable example of an invasive species. It has been well documented that the population growth and the range of expansion of these birds coincide with the urbanization process (Arnold & Folse, 1977; Dinsmore & Dinsmore, 1993; Peer, 2011; Pratt, Ortego, & Guillory, 1977; Pruitt & McGowan, 1975; Wehtje, 2003). Casual observations suggest that grackles benefit from the resources of human activities, such as the frequent provision of food and nesting materials from trash, which probably helps this species of bird to succeed in a wide range of ecological niches in the American continent. Nevertheless, there is little evidence about its feeding patterns and social interactions during the day that can support such claims (K. Johnson & Peer, 2001).

Two models of predation-starvation risk in birds predict how the foraging behavior and the frequency of animals should change throughout the day. The bimodal foraging hypothesis, predicts (as his name states) a bimodal relation between the foraging intensity and the time of the day in birds living in environments with stable access to food and quickly replenishable energy reserves (McNamara, Houston, & Lima, 1994). The assumption is that during morning the birds recover some of the weight they lose as a result of the absence of food consumption during the previous night, followed by an abrupt decrease in activity during afternoon to save energy reserves and avoid predators, and finally, an increment on activity to gain reserves for the upcoming night. In this model, the excessive gain of fat could affect the maneuverability and speed of flight, which can turn a bird into a more prone prey. On the other hand, the risk-spreading theorem specifies that if the risk of predation does not alter the daily foraging patterns of the birds or this factor has minimal influence, then it would be expected a constant foraging activity throughout the day (Houston, McNamara, & Hutchinson, 1993). In this way, observe the behavior of grackles in a patch with low and high resources availability will allowed us to determine which model could explain better the behavior of the grackle. Also, the Marginal Value Theorem (MVT; Charnov, 1976) applied to our case suggest that a higher cost of exploitation should promote lower number of subjects in the patch because the optimal strategy would be to leave the site and look for another place in a rich trash-environment, like the city. So, in a situation with different degrees of resources availability we should find also different frequency of subjects exploiting the resources.

If the grackle is a social species, we should observe differences in their interaction according to the hierarchy levels, so the members of the group could play the role of scrounger or producer according to such level but also this interaction could evidence if the specie do group foraging. Finally, if grackles are social animals, we should observe consistent relations between their foraging activities, the subjects foraging, and their social interactions. As a first approach to the problem we located a garbage patch outside a residential building, in a high-density urban area, where the residents left their trash for pickup and a group of wild Great-tailed Grackles (*Quiscalus mexicanus*) were identified as regular visitors. The main aim of this research was to differentiate if the daily foraging patterns of grackles depend on the availability of the resources, by recognizing if the strategy used in foraging changes as a function of the presence or absence of garbage collection. At the same time, it was intended to know if the interactions between the members of the group were determined by the availability of food at the patch during the day, through correlating the number of grackles present at the observation site and the number of aggressive behaviors. Furthermore, it was also explored if the hierarchy plays a role in the agonistic behaviors, taking into consideration the moment of arrival of the subjects to the feeding patch. At last, an attempt was made to establish if social behavior affects the capacities of these birds to survive hazardous environments such as cities.

## Method

### Subjects and observation site

We locate a territory range of four trees near a residential building in a high-density urban area in the vicinity of the metropolitan area of Guadalajara, Jalisco (México) where a group of at least eight wild Great-tailed Grackles (*Quiscalus mexicanus*) lived. During the observation, it was not possible to identify each subject.

This site was chosen due to the regular visits of grackles in the area most likely as an effect of the recurrence with which residents left their trash for pickup. The approximate area of the observation site was of 50 m^2^ (10 meters long by 5 meters’ width). At the center of the observation site, there was a concrete tree pot (approximately 200 cm long by 200 cm width and 70 cm tall), with a sown pine at the right side of it. On the left side of the pot, there were two garbage bins (made of oil barrels, which can hold approximately 159 liters). The residents adopted the left surface of the pot as a secondary place to leave their trash.

Because the observations were carried out in natural conditions, the observers sometimes saw people leaving or digging the garbage, parking their cars, sweeping the street or sometimes walking their pets. In addition to this, the arrival of animals such as dogs, cats, pigeons, and finches was witnessed at the observation site. None of these events were recorded.

### Video Recordings

The recordings were carried on between 0630- and 2200-hours during November 2017 and January 2018. A Sony Handycam® DCR-SR85 was placed inside one of the apartments in the residential area, approximately 15 meters above street level. The camera was always pointing towards the garbage bins and the concrete pot. A total of 194 hours extracted from 25 days of recordings were analyzed. In twelve of the 25 videos, the collector truck passes by six times on Wednesday, four on Friday, one on Monday and one on Saturday. The remaining 13 videos were recorded in the absence of the collector truck, and four occurred on Thursday, three on Tuesday, three on Saturday, two on Sunday and one on Monday.

### Observers’ Training

Three observers recorded the videos. Before starting the recording, we trained the observers by selecting video fragments of 30 minutes randomly. These portions of the video were delivered to the observers independently, and they were asked to record the behaviors of the ethogram (see Table 1).

In order for the observers to keep an adequate record of the events, the following rules were established: a) the start of the grackle registration began when the first subject (from now on referred as focal subject) appeared complete or at least 50% of his body, including head and half of the back; b) the registration of the focal subject ended when the grackle left the shot or if an object blocked the visualization of the bird for more than 3 seconds; c) if the subjects arrived at the study area flying, the registration was initiated only when they prostrated at any angle of the shot that allowed their registration; and, d) if the grackles flew by and did not prostrate somewhere, there was no activity recorded.

Once the registration was completed, we calculate the reliability among the observers. If the coefficient of inter-observer reliability was below 0.7, the errors were reviewed, and the registration criterion was agreed upon and reconciled. A new fragment of video of 30 minutes was delivered, and a new observation was carried out.

Finally, when the level of reliability between observers reached approximately 0.8, a final test was made with a 3-hour video. After the observers reached the desired level of reliability, the 25 videos were randomly divided.

All observers watched eight videos each, a ninth video was analyzed by the three of them, and with that recording, the Cohen’s κ was run to determine if the coefficient of inter-observer reliability of the analyzed videos was accomplished. There was a substantial agreement between the observers, κ=.792 (Landis & Koch, 1977). Approximately every observer examined 65 hours.

### Ethogram and registered events

During the first phase of the observers training, we established the ethogram to be analyzed. Most of the behaviors included in the ethogram were aimed at analyzing the behavior of the focal subject (i.e., at the observation site, fly, searching for food, jump, displacement, out of the site). While others were intended to mark the occurrence of other events such as the arrival of more subjects to the location or the garbage collection (see Table 1).

It is important to emphasize that the focal subject was the first grackle to arrive at the observation site. Upon arrival, all focal behaviors were recorded. Once the focal subject left the observation site, only the number of subjects present at the observation site or garbage collection events were recorded, if it was the case. The registration of a new focal subject could not occur again until all the grackles of the group abandoned the observation site. In such a way, any member of the group could be considered as a focal subject, if he was the first to arrive at the observation site.

The behaviors defined in the ethogram as states (behaviors with appreciable durations) were *foraging, at the observation site, and out of the site*. Those defined as events (behaviors that occurred at an instant) were *fly, jump, aggression, garbage*, and the number of grackles (i.e., 1 thru 8 or more; Altmann, 1974). There was an exclusion criterion between behaviors to avoid errors during the registration. *Fly, jump, aggression*, and *out of the site* stopped the *foraging* behavior registration automatically; and *foraging* could not have occurred when the grackle was *out of the site*.

The absence of garbage collection refers to those days when the garbage truck did not collect the garbage, and therefore, the waste remained inside the plastic garbage bags. While the events in presence, refer to those days when the truck collected the trash and part of the organic and inorganic waste was scattered on the ground.

### Data Analysis

We measure the frequency of foraging behaviors, the number of subjects present in a feeding site, and the frequency of aggression between individuals. The analysis of the videos was carried out with the BORIS registration software (v.6.0.6) installed on computers with Windows operating system (Friard & Gamba, 2016). We used the coefficient of inter-rater reliability tool included in the software to calculate Cohen’s kappa. For this analysis we excluded the behaviors *fly, jump, and out of the site* and we used time slides of 0.5 seconds. Once the event registration was completed, a data matrix was created and analyzed with MATLAB (v. R2019a). To detect possible associations between the time of the day and the independent variables measured (i.e., foraging, and aggressions) we calculate the area under the curve (AUC; Pruessner, Kirschbaum, Meinlschmidt, & Hellhammer, 2003) with the formula:

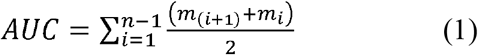

Here *m_i_* is each value of the independent variable and *n* the total amount of measurements. We decide to use this measure to compare if there were variations in the distribution of foraging and aggression, depending on the presence or absence of garbage collection throughout the day. During the results, three different ways were used to describe the AUC data in each of the registered variables. The first represents the AUC of the total recorded events (Total AUC) calculated with formula (1). The second, the percentage of the total AUC (% AUC) and is obtained by dividing each AUC event by the total AUC and multiplying the result per 100. The third indicates the percentage of AUC events for each interval of the day (% AUC per interval), and it is calculated by dividing each AUC event among the total AUC of a specific interval (i.e., 7 to 11, 11 to 15 or 15 to 20) and multiplying the result by 100. The tables shown in the results only show the total AUC, and the percentage of AUC.

## Results

The distribution of foraging in the presence of garbage collection is like a normal distribution but in its absence has several peaks (Fig. 1A). The percentage of area under the curve showed that 15.35% of the total events recorded occurred in the morning (0700 – 1100 h); while at noon (1100 – 1500 h) the grackles produced up to 57.10% of these behaviors; and finally, during the afternoon and evening (1500 -2000 h) the foraging decreased to 27.55% (Table 2).

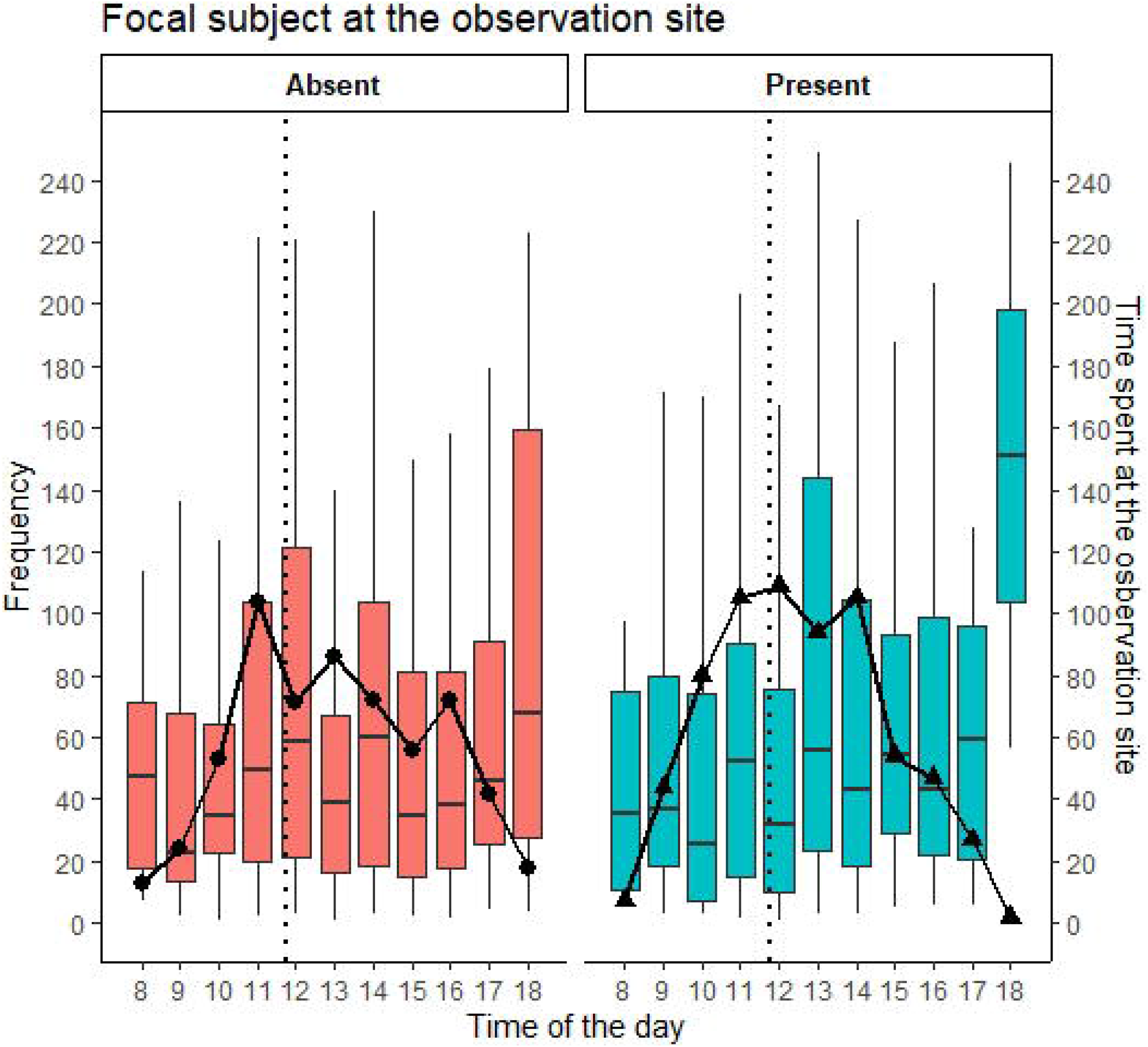

In total, the foraging registered in the presence of garbage collection was 4.89% higher than in its absence (see Table 2). In both cases, the distribution of foraging begins at 0800 h; however, the percentage of the area under the curve per interval from 0700 to 1100 h shows that the growth of the distribution in the presence of garbage collection was 18.39% more than in its absence (Table 2). The frequency peak of foraging behaviors in the presence of garbage collection occurred at 1300 h but remained relatively constant since 1100 h up until 1400 h 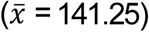. Conversely, foraging in the same period decreased in the absence of garbage collection 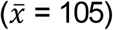, for this distribution, the maximum peak occurred at 1200 h. It is interesting to note, that the start of the stability period of foraging in the presence of the collector truck coincides with the maximum peak of foraging those days (black dashed line, Fig. 1A) that occurred at 1100 h, although the latter could vary between 1000 and 1400 h over the days. The percentage of AUC during noon (1100 – 1500 h) shows that the frequency of foraging in the presence of garbage collection was 13.48% more than in its absence (Table 2).

In both foraging frequency distributions, there is a sudden decrease in grackles activity at 1500 h which slightly increased towards 1600 h. The foraging at 1600 h in the absence of garbage collection was 114, less than in its presence 77, a difference of 19.37%. Finally, foraging decreased progressively in both distributions (1500 – 2000 h). In the presence of garbage collection, the foraging behavior ended at 1800 h, but in the absence of garbage collection, it extended up until 1900 h, the percentage of AUC per interval shows an increase of 20.43% (Table 2). After this time, grackles ceased their activity.

At all times the number of grackles present at the observation site was more during the presence of garbage collections than in its absence, increasing gradually up until 1500 h and diminishing after that (Table 3). The total events recorded when more than one grackle was at the observation site was 6756 events. The 57.13% of the interactions (events with more than one grackle) during the day occurred in the presence of garbage collection (Figure 1B) while the 42.87% of them happened in its absence; a difference of 14.27% (Table 3).

Records of interactions between two to four grackles were 15.66% more frequent 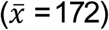 in the presence of garbage collection between 1100 and 1400 h than in its absence 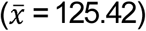. The highest frequency of recorded interactions in both conditions was between two subjects 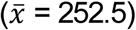 at 1200 and 1300 h, which represents the 3.74% of the total registered events. The interactions between grackles in the presence of garbage collection increased gradually after 0900 h, staying relatively constant 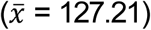 between 1100 and 1700 hours and reducing their frequency towards 1800 h. Although this was similar for interactions in the absence of garbage collection events, the frequency of events was 14.69% less throughout the day 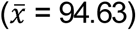. The interactions between six or more grackles at the observation site represents 7.65% of the total events of which, 5.17% occurred the days of garbage collection while 2.49% happened other days, which represents 35.01% more interactions in the former than the latter, although these interactions were infrequent.

More aggressive behaviors in the absence of garbage collection, 58.82%, than in presence, 41.18%, were recorded, a difference of 17.65%. The same was observed at different hours, the percentage of area under the curve between 0700 and 1100 h shows 24.14% more aggressions in absence than in the presence of garbage collection, 34.57% from 1100 to 1500 h, and 39.73% at 1500 to 2000 h. So, even when the frequency of activity was low at the beginning and end of the day, the aggressions were more frequent the days in absence of garbage collection, with low resources availability (see Figure 2).

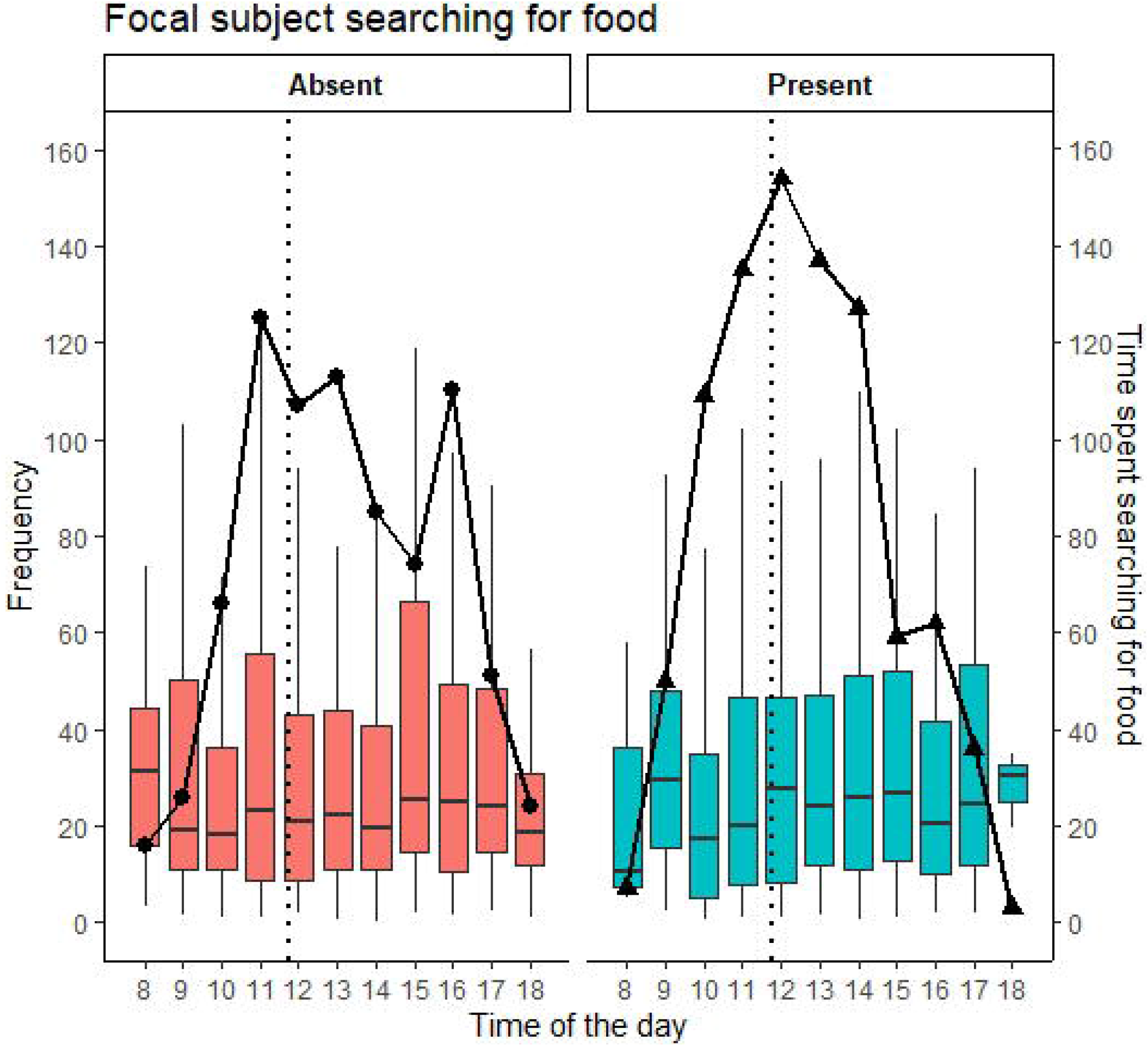

It can be noted that the percentage of the area under the curved show that the 31.82% of aggressive behavior occurred in the presence of garbage collection, of which, 12.50% of these behaviors were from the focal subject to others, and 19.32% were from other subjects to the focal. Likewise, the 68.18% of aggressive behaviors happened in the absence of garbage collection, 22.73% of them were from the focal subject to others, and the 45.45% were from others to the focal (Table 4). Most time the aggressions were in the direction of other subjects to the focal, and such frequency was higher the days without garbage collection.

Similarly, at analyzing the results of the total area under the curve (Table 4), the aggressive behaviors between 1100 to 1500 h from the focal subject to others (9.5%) was 25.49% less than the aggressive behaviors from other subjects to the focal (16%) in the presence of garbage collection (25.5%). The difference was similar between 0700 to 1100 h and 1500 to 2000h, being respectively 14.29% and 15.38%. On the other hand, the total area under the curve for the aggressions from other subjects to the focal in the absence of garbage collection were similar to those described above between 1100 to 1500 h (38.74%) and 1500 to 2000 h (53.19%), but from 0700 to 1100 h the displacement events where 36.36% more frequent from the focal subject to others.

The relation between the frequency of aggression and the frequency of grackles at the observation site was positive as should be expected, but the frequency of aggression to the focal is notably higher the day without garbage collection (See Annex, Figure 1). The correlation between foraging frequency and events of more than one grackle at the observation site in the presence (solid black line) and the absence (solid gray line) of garbage collection (Figure 3) shows that the higher the foraging behavior, the higher the frequency of more than one grackle in both conditions. The relation is a straight line in both situations. It is important to remark that each point represents the time of the day when the observers recorded the events.

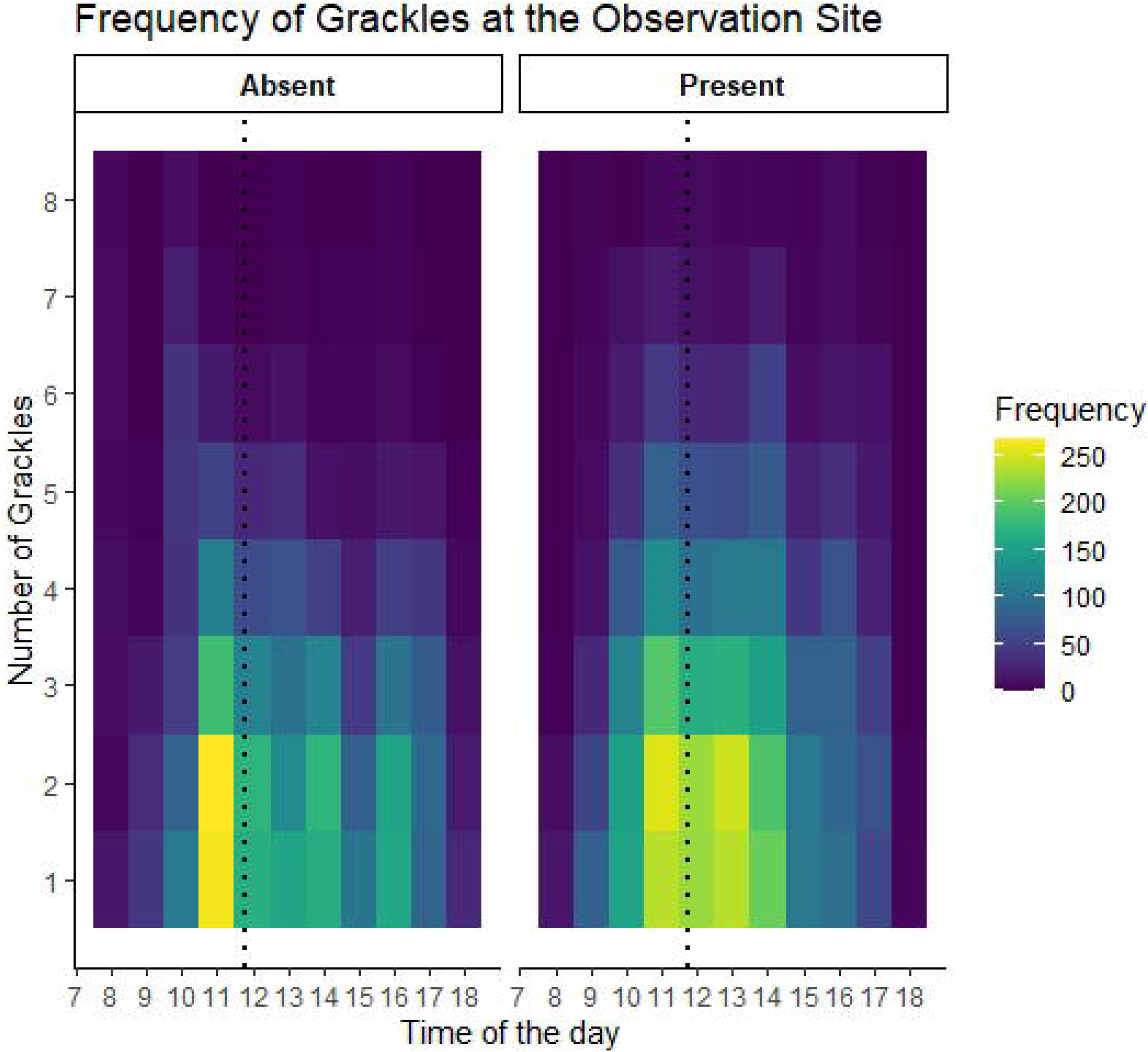

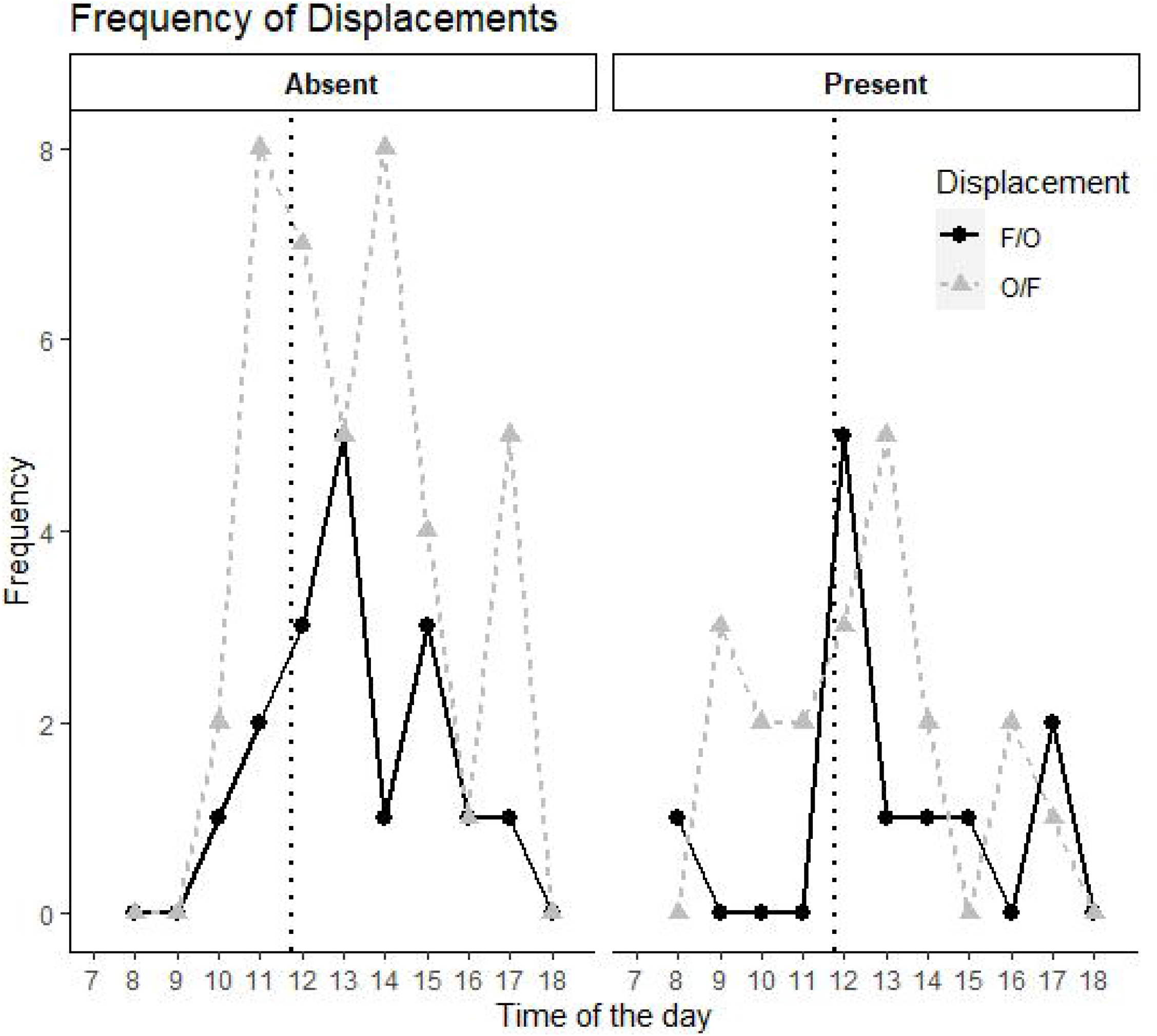

There are slight differences between the parameters of the linear regressions in the presence (y= 0.601x + 1.61, r^2^ = 0.98) and absence (y= 0.599x + 3.13, r^2^ = 0.947) of garbage collection. The functions grow with similar slopes, but the intercept is closer to zero the days with garbage collection, meanwhile, in the absence, there is more activity represented by a higher intercept. In the absence of garbage collection, the animals seem slightly more active, but their foraging activity increased with the number of subjects at the site for both conditions. However, it can be observed that in the presence of garbage collection, the frequency of recordings of more than one grackle was more frequent between 1100 and 1400 h compared to those days in the absence of garbage collection, and the same happened with foraging.

It is interesting to note that the function between the foraging frequency and the number of grackles at the observation site is a straight line with positive slope; this suggests that grackles are a species of birds that feed as a group. The foraging frequency is the activity of the focal subject, and this subject is more active when more grackles are at the site. However, both things occur at the same time of the day so the relation may be result of higher levels of activity during noon. Notwithstanding the focal subjects could abandon the place due to the presence of the others, the animals continue and increase foraging, so the presence of other subjects does not interfere with the foraging, if the presence of other subjects has an effect is more in the direction of incentive behavior. Nevertheless, an experimental separation of the variables (i.e., time of day and number of subjects) would be necessary to establish the effects.

## Discussion

The present research aimed to answer the question of whether the grackles can obtain resources of biological relevance from human waste, by analyzing if the foraging and aggressive behaviors were modified when garbage collection occurred or not. Mostly the results showed that garbage collection (availability of resources) is associated with a higher frequency of foraging, social interactions, and a higher number of subjects at the site. In the presence of garbage collection, foraging, and the number of individuals at the observation site was higher than in its absence. Conversely, the number of aggressions was higher in the absence of garbage collection than in its presence.

The specific goals of this work have focused on answering if the availability of resources influenced the foraging activity of grackles, their social behaviors, the number of subjects present at the observation site and the aggressive behaviors.

The first question was aimed at knowing if the availability of resources modified the grackles’ daily foraging patterns. The results showed that grackles spend more time searching for food during noon than morning and evening. The main difference between conditions of resource availability is that in the presence of garbage, the daily foraging patterns show a higher frequency throughout the day which assembly a normal distribution, opposite to those days in absence where the pattern has several peaks and lower frequency (Table 2 and Figure 1A).

As mentioned before, in the bimodal foraging hypothesis, the birds should limit their behavior to dawn and dusk to avoid the risk of predation during noon but also prevent starvation. Contrarily, the risk-spreading theorem states that if the predation does not influence the foraging behaviors, then the birds should forage constantly throughout the day. Our results on the frequency of foraging in the absence of garbage collection seem to follow the pattern suggested by the bimodal foraging hypothesis, increased activity during the morning followed by a long paused and an increment at the end of the day (dotted line, Figure 1A). However, the distribution of foraging when the garbage collection happened gives support to the risk-spreading theorem, because grackles seem to have higher activity during the morning and a steady response pattern during noon (black line, Figure 1). The latter foraging distribution is similar to Bonter, Zuckerberg, Sedgwick, and Hochachka, (2013) data. In this experiment, the researchers show that the rate of visits to the feeder during winter of four different species of Passeriformes (*Poecile atricapillus, Baeolophus bicolor, Sitta carolinensis* and *Haemorhous mexicanus*) supports the predictions of the risk-spreading theorem and reject the bimodal pattern. Their results show the rate of visits increase hour by hour reaching a maximum peak 2 hours before sunset and an accelerated decrement before nightfall.

Although the foraging distribution in the absence and presence of garbage collection seem to support different models, they could be explained as an effect of the short-term plasticity that grackles could have developed to survive hazardous environments such as cities. Throughout the observations, we observed that grackles could obtain food in two different ways; by tearing the plastic bags inside the garbage bins (which most –if not all-times were present) looking for organic waste or insects at the bottom of them or by getting the food with the help of sanitation workers.

In Mexico, it is common that part of the organic and inorganic waste kept dispersed in the ground for an indefinite period at the end of the garbage collection. During this time, it seems that grackles take advantage to obtain the food spread. It appears that getting resources from garbage requires a higher cost of response in the absence (i.e., tear plastic bags) and a lower cost of response in the presence of garbage collection (i.e., directly foraging from the ground).

As we have mentioned before, most garbage collection happened on Wednesdays and Fridays, while most of the recordings in the absence occurred on Tuesdays, Thursdays, and Saturdays. That means every day, the availability of resources is different but not as an effect of absolute lack of resources, but as an effect on the difficulty to obtain it. In other words, grackles have a more challenging task to get the food when the garbage is still packed at plastic bags that when the sanitation worker disperses the food when recollecting the garbage. The latter can lead grackles to leave the food patch in days in which the cost of getting the food is high, and the starvation is raised than in those days in which it was easier to get it. In this way, the decrease in foraging the days without resource availability (without garbage collection) can be explained as an effect of prolonged deprivation but not as an effect of saving energy resources or as a strategy to avoid predators just as in the bimodal hypothesis.

Additionally, it could be possible that the decreased foraging activity during the evening in those days, in which the collector truck was present, is the result of a possible satiety effect derived from the constant feeding activity during noon. If satiation is high, then the animals stop foraging; otherwise, the behavior sustains until dawn. In those days in which grackles are already satiated, they cease foraging earlier than when they are deprived. All these elements together seem to support our hypothesis that the garbage collection increased the availability of resources as a result of the dispersion of food residues in the ground, which facilitated the consumption. In other words, the easiest way to obtain the resources, the higher would be the time spent in obtaining them.

In addition to the above, we believe it is important to highlight two events that occurred during foraging. First, at the beginning of the day, the activity in the presence of garbage collection increased more rapidly than in its absence. This effect made us think if grackles were able to anticipate the arrival of the collector truck. Although we cannot assure if these birds can anticipate, we think grackles can discriminate events such as the arrival of residents to leave their trash, which prone grackles to start the foraging early in the morning.

Second, the decrement observed in both foraging distributions between 1500 -1600 h could be due to an increase in human activity near the observation site. The observations lead us to note that the residents tend to arrive at their homes at this interval which coincides with the lunch hour of the citizens who work in the area and the time the residents take to walk their pets. Although we did not register the arrival of another species, we notice that when mammal species (including humans, dogs or cats) enter the site most of the times the group of grackles leaves the observation site, but also, we recognize that grackles could share the same area with other bird species. These results match with the descriptions made by Johnson and Peer, (2001). The authors have registered that grackles share their territory with species such as Purple Gallinule (*Porphyrio martinica*), American Coot (*Fulica americana*), Common Moorhen (*Gallinula galeata*), Pied-billed Grebe (*Podilymbus podiceps*), Least Grebe (*Tachybaptus dominicus*), and Green Heron (*Butorides virescens*) and also other blackbird species.

The second question addressed was to recognize if the availability of food affects the social interactions between the members of the group, especially if the number of grackles at the patch during foraging, change depending on the presence or absence of garbage collection. Likewise foraging, the results always showed that the number of grackles was higher in the presence of garbage collection than in its absence. Also, the most commons interactions were between two to four subjects, the higher one between two subjects, while the interactions of six or more subjects were infrequent (Table 3, and Figure 1B). Apparently, the lower cost of obtaining resources associated with the dispersion also attracts a higher quantity of grackles at the observation site. Whereas, in the days when the garbage collection did not occur, the number of grackles decreased as an effect of the deprivation, which in turn favored the abandonment of the patches.

According to the above, it seems that grackles prefer to spend the least amount of energy tearing the bags and with it, a shorter time in obtaining the food than to go in search of other patches, then our data follow the marginal value theorem. Thus, as stated by Charnov, (1976) there is a relation between the cost of response associated with obtaining food and the abandonment of foraging patches by the members of the group. Therefore, possibly the number of grackles at the observation site in the presence and absence of garbage collection is linked to the cost of obtaining food and the state of deprivation. Though, the constant availability of resources could explain the growth and establishment of grackles in urban areas. According to Pyke, Pulliam, and Charnov, (1977), the less variability exists in the food availability of the patches present in the environment, the likelihood that the animals seek alternatives will be less, which could promote the settlement of the species.

On the other hand, when correlating the number of grackles present at the observation site and the foraging, we found similar slopes and positive increases in the absence and presence of garbage collection; which confirm an intimate relation between the variables (Figure 3). This result suggests that this species of bird is highly social and possibly the mechanisms used to obtain resources and exploitation of the environment is regulated by the interaction between the individuals of the group, which could explain one of the reasons of success in this species even in hostile environments such as cities. Even though, it is necessary to develop a study that can deepen more about the social relation among grackles and its effects on adaptation to the cities.

On our third question, we intend to identify if the garbage collection altered the display of agonistic behaviors and if the hierarchy played a role. The results showed there was more aggression in the absence of garbage collection than in its presence. Consistently, the number of aggressions from the focal animal (the first subject to arrive) to the other subjects was less frequent than from others to the focal in both conditions (Table 4, and Figure 2). Moreover, we found a positive correlation between the aggression and the number of grackles; and persistently, the slopes were higher in the absence of garbage collection than in the presence (Figure 1, annexes).

As has been described in the producer-scrounger game literature, in assessing dominance as a reason for agonist behaviors, the dominant subjects parasitize more the less dominant individuals (Barta & Giraldeau, 1998). The results of the present study (Figure 2) show that the focal animal tends to exhibit a lower number of aggressive behaviors towards the other birds of the group and vice versa, the other subjects of the group must display a higher number of aggressive behaviors towards the focal animal. If we consider that the focal animal points out the source of food and that the other subjects arrive to parasitize, then, we suppose that, for grackles, the agonist behaviors are mean to obtain resources. The establishment of dominance or hierarchy could be like that reported in other animal species. For example, female baboons scrounge more on subordinate individuals whit lower ranks (King, Isaac, & Cowlishaw, 2009), rooks which show more dominance by chasing and pulling feathers of other members of the group (Jolles, Ostojić, & Clayton, 2013), and biggest salmons’ which strikes the smallest ones (Phillips et al., 2018). Of course, to confirm these assumptions it is necessary to extend the current results, maybe by identifying the subjects observed.

Although we did not register the sex of the individuals, the observers underline (based on phenotypic differences) that males made most of the aggressive behaviors. If the scrounger animal was the dominant male within the group, our results are similar to some reports that show that dominant grackles exhibited most of the agonistic behaviors (i.e., the number of movements towards an opponent) towards trespassing rival males, lower ranked males, females, and juveniles to maintain stable dominance relations within the group (K. Johnson & Peer, 2001).

It is essential to point out that the higher number of aggressions of the focal subject to others during the period from 0700 to 1100 h may be due to a prolonged deprivation effect. In this situation, it seems that the focal subject decides to remain for even longer, even in the face of aggressions from other subjects, which results in a higher defense of resources. However, with the arrival of more subjects as time passes, the focal subject tends to withdraw.

By our results, we can conclude that grackles can adapt to slightly changes in the environment that occurred day by day, such as the presence or absence of the garbage collector truck. We also can claim that the continuous garbage waste that occurs in cities is a reliable source of energy, and the ease in obtaining it can play an essential role in their social lives, such as changes in the foraging or the synergy within a group.

Most importantly, this research open the possibility to explore questions about the cognitive abilities of these birds by placing restrictions in the environment — questions more attached to the experimental research than the observational one (for some advances in the area see, Logan, 2016a, 2016b, 2016c) . Questions that can help researchers explore how the cognitive skills of these birds can help them thrive in such a changing environment as cities.

## References

Altmann, J. (1974). Observational Study of Behavior: Sampling Methods. Behaviour, 49 (3), 227–266. https://doi.org/https://doi.org/10.1163/156853974X00534

Arnold, K. A., & Folse, L. J. (1977). Movements of the Great-Tailed Grackle in Texas. The Wilson Bulletin, 89 (4), 602–608. Retrieved from http://www.jstor.org/stable/4160987

Barta, Z., & Giraldeau, L. A. (1998). The effect of dominance hierarchy on the use of alternative foraging tactics: A phenotype-limited producing-scrounging game. Behavioral Ecology and Sociobiology, 42 (3), 217–223. https://doi.org/10.1007/s002650050433

Bonter, D. N., Zuckerberg, B., Sedgwick, C. W., & Hochachka, W. M. (2013). Daily foraging patterns in free-living birds: Exploring the predation-starvation trade-off. Proceedings of the Royal Society B: Biological Sciences, 280 (1760), 1–7. https://doi.org/10.1098/rspb.2012.3087

Charnov, E. L. (1976). Optimal Foraging, the Marginal Value Theorem. Theoretical Population Biology, 9(2). https://doi.org/10.1016/0040-5809(76)90040-x

Dinsmore, J. J., & Dinsmore, S. J. (1993). Range Expansion of the Great-tailed Grackle in the 1900s. Journal of the Iowa Academy of Science, 100 (2), 54–59. Retrieved from https://scholarworks.uni.edu/jias/vol100/iss2/4

Evans, K. L., Newton, J., Gaston, K. J., Sharp, S. P., McGowan, A., & Hatchwell, B. J. (2012). Colonisation of urban environments is associated with reduced migratory behaviour, facilitating divergence from ancestral populations. Oikos, 121 (4), 634–640. https://doi.org/10.1111/j.1600-0706.2011.19722.x

Friard, O., & Gamba, M. (2016). BORIS: a free, versatile open-source event-logging software for video/audio coding and live observations. Methods in Ecology and Evolution, 7 (11), 1325–1330. https://doi.org/10.1111/2041-210X.12584

Gallo, T., & Fidino, M. (2018). Making wildlife welcome in urban areas. ELife, 7. https://doi.org/10.7554/eLife.41348

Houston, A. I., McNamara, J. M., & Hutchinson, J. M.C. (1993). General results concerning the trade-off between gaining energy and avoiding predation. Philosophical Transactions of the Royal Society of London. Series B: Biological Sciences, 341(1298). https://doi.org/10.1098/rstb.1993.0123

Johnson, K., & Peer, B. D. (2001). Great-tailed Grackle (Quiscalus mexicanus), version 2.0. In A. F. Poole & F. B. Gill (Eds.), The Birds of North America. Retrieved from https://doi.org/10.2173/bna.576

Johnson, M. T. J., & Munshi-South, J. (2017). Evolution of life in urban environments. Science, 358 (6363), eaam8327. https://doi.org/10.1126/science.aam8327

Jolles, J. W., Ostojić, L., & Clayton, N. S. (2013). Dominance, pair bonds and boldness determine social-foraging tactics in rooks, Corvus frugilegus. Animal Behaviour, 85 (6), 1261–1269. https://doi.org/10.1016/j.anbehav.2013.03.013

King, A. J., Isaac, N. J. B., & Cowlishaw, G. (2009). Ecological, social, and reproductive factors shape producer-scrounger dynamics in baboons. Behavioral Ecology, 20(5). https://doi.org/10.1093/beheco/arp095

Landis, J. R., & Koch, G. G. (1977). The measurement of observer agreement for categorical data. Biometrics, 33 (1), 159–174.

Logan, C. J. (2016a). Behavioral flexibility and problem solving in an invasive bird. PeerJ, 4:e1975. https://doi.org/10.7717/peerj.1975

Logan, C. J. (2016b). Behavioral flexibility in an invasive bird is independent of other behaviors. PeerJ, 4:e2215. https://doi.org/10.7717/peerj.2215

Logan, C. J. (2016c). How far will a behaviourally flexible invasive bird go to innovate? Royal Society Open Science, 3 (6), 1–10. https://doi.org/10.1098/rsos.160247

McKinney, M. L. (2002). Urbanization, Biodiversity, and Conservation: The impacts of urbanization on native species are poorly studied, but educating a highly urbanized human population about these impacts can greatly improve species conservation in all ecosystems. BioScience, 52 (10), 883–890. https://doi.org/10.1641/0006-3568(2002)052[0883:UBAC]2.0.CO;2

McNamara, J. M., Houston, A. I., & Lima, S. L. (1994). Foraging Routines of Small Birds in Winter: A Theoretical Investigation. Journal of Avian Biology, 25 (4), 287–302. https://doi.org/10.2307/3677276

Peer, B. D. (2011). Invasion of the Emperor’s Grackle. Ardeola, 58 (2), 405–409. Retrieved from https://doi.org/10.13157/arla.58.2.2011.405

Phillips, J. A., Peacock, S. J., Bateman, A., Bartlett, M., Lewis, M. A., & Krkošek, M. (2018). An asymmetric producer-scrounger game: body size and the social foraging behavior of coho salmon. Theoretical Ecology, 11 (4), 417–431. https://doi.org/10.1007/s12080-018-0375-2

Pratt, H. D., Ortego, B., & Guillory, H. D. (1977). Spread of the Great-Tailed Grackle in Southwestern Louisiana. The Wilson Bulletin, 89 (3), 483–485. Retrieved from http://www.jstor.org/stable/4160959

Pruessner, J. C., Kirschbaum, C., Meinlschmidt, G., & Hellhammer, D. H. (2003). Two formulas for computation of the area under the curve represent measures of total hormone concentration versus time-dependent change (multiple letters). Psychoneuroendocrinology, 28(7), 916–931. https://doi.org/10.1016/j.psyneuen.2003.10.002

Pruitt, J., & McGowan, N. (1975). The return of the great-tailed grackle. Amer. Birds, 29, 985–992.

Pyke, G. H., Pulliam, H. R., & Charnov, E. L. (1977). Optimal Foraging: A Selective Review of Theory and Tests. The Quarterly Review of Biology, 52 (2), 137–154. https://doi.org/10.1086/409852

Sol, D., Lapiedra, O., & González-Lagos, C. (2013). Behavioural adjustments for a life in the city. Animal Behaviour. https://doi.org/10.1016/j.anbehav.2013.01.023

Wehtje, W. (2003). The range expansion of the great-tailed grackle (Quiscalus mexicanus Gmelin) in North America since 1880. Journal of Biogeography, 30 (10), 1593–1607. https://doi.org/10.1046/j.1365-2699.2003.00970.x

